# Endoluminal catheter pulsed field ablation for the treatment of atherosclerotic vascular disease

**DOI:** 10.1101/2025.09.06.674641

**Authors:** Zaid Salameh, Edward Jacobs, Rafael Davalos

## Abstract

**Background:** Atherosclerotic vascular disease remains a leading cause of morbidity and mortality worldwide. Current treatments such as angioplasty, stenting, and atherectomy are invasive and limited by restenosis, thrombosis, and incomplete long-term efficacy. Pulsed field ablation (PFA), a nonthermal electroporation-based modality, has demonstrated safety in other cardiovascular applications, but it has not been applied for the treatment of endoluminal vascular diseases. We investigated whether pulsed electric fields could be delivered within the coronary artery and if PFA could selectively ablate the cellular components of atherosclerotic plaques.

**Methods:** A bipolar catheter-based PFA probe was fabricated using a balloon catheter with flexible electrodes and evaluated through a combination of *ex vivo* evaluations. The electrical conductivities of human atherosclerotic plaques were derived from previous impedance measurements for patient-specific multi-tissue and single-cell electroporation modeling. PFA was then evaluated for selective decellularization within an electrical conductivity-matched 3D fibrotic atherosclerosis tissue mimic using high concentrations of human macrophages and aggregated oxidized low-density lipoproteins, encapsulated within a collagen matrix.

**Results:** Endoluminal bipolar probe evaluation demonstrated probe positioning and high voltage pulsed electric field delivery within the left coronary artery of *ex vivo* porcine hearts, with maximum ablations (6.99 cm^2^) and current (13 A) evaluated within live potato tissue. The multi-tissue model then indicated that endoluminal PFA can effectively cover >95% of severe and thick plaques with irreversible electroporation, with single-cell modeling supporting the electroporation of foam cells within the plaque. The 3D atherosclerosis mimic validated the ability of PFA to completely ablate the foam cells with fibrotic tissue at >1000 V/cm.

**Conclusions:** This study provides the first demonstration of PFA for the treatment of atherosclerotic vascular disease. By combining experimental validation with computational modeling, we establish proof-of-concept that PFA can selectively ablate diseased cells while preserving extracellular architecture, laying the groundwork for future translational development of this therapy.

## Introduction

Vascular diseases (VDs) are the most severe global health concern, with approximately 17.9 million yearly deaths (∼32% of all global deaths), and are the leading cause of mortality within the United States^1^. The number of patients diagnosed with VDs continues to increase yearly due to worsening lifestyles and an aging population, and many common chronic conditions, such as hypertension and diabetes, are closely linked to VDs^2^. Vascular dysfunction can lead to reduced blood flow, tissue ischemia, and organ damage, ultimately contributing to serious health outcomes like stroke, heart attack, limb amputation, and kidney failure. Specifically, intravascular conditions that involve the inner lumen or walls of blood vessels play a central role in disease progression. Intravascular abnormalities such as atherosclerotic (AS) plaque formation, thrombosis, and inflammation can disrupt normal blood flow, trigger clot formation, and lead to vascular occlusion or rupture^3,4^.

AS is the primary source of vascular dysfunction, developing from plaque formation within the intimal layer of arteries^5^. Typically, AS is initiated by the passage of low-density lipoprotein cholesterol (LDL-C) through the arterial endothelium, which accumulates within the intima. The LDL-C induces the upregulation of adhesion molecule expression on the vascular endothelium. An abnormal and turbulent flow within the arterial lumen can also elevate the risk of AS by concomitantly promoting the expression of adhesive molecules, with plaque narrowing further exacerbating the condition^6^. The progression of AS depends on the activation of the inflammatory response, whereby monocytes infiltrate the lesion site, differentiate into macrophages, and internalize oxidized cholesterol, ultimately differentiating into foam cells^7^. Foam cells are the primary cellular components found in the plaque and may locally proliferate to advance the disease^8^. The smooth muscle cells (SMCs) within the blood vessel are also crucial in progressing the disease, as they can also internalize oxidized cholesterol to become foam cells and upregulate macrophage and endothelial markers, contributing to plaque destabilization^9^. If a too large of a plaque forms, then apoptotic cells in the center may not be removed, leading to a necrotic core that impedes the distribution of targeted therapies into the plaque^10^. Destabilized plaques often occur at advanced stages of AS, which can rupture and lead to thrombosis. It is crucial to stabilize or remove plaques before thrombosis formation, as they can block downstream vasculature, leading to Major Adverse Cardiovascular Events (MACE), such as myocardial infarction or stroke. Most AS is diagnosed at these late stages, necessitating immediate treatment to prevent potential MACE.

The biological processes outlined are not only central to the pathology of AS development but also represent key targets for current therapeutic intervention. Current therapeutics for AS typically use statins or anti-inflammatory drugs to inhibit foam cell proliferation, thereby stabilizing the plaque and potentially preventing a MACE. However, these therapeutics do not reverse plaque formation, with many patients experiencing intolerance to long-term therapy. Stents are a common long-term solution, but they do not remove the plaques; instead, they reshape and compress them to open the vessel and improve blood flow. Stents do not prevent vessel hardening, which can lead to hemorrhaging, and still possess a risk for plaque development and rupture^11^. If the plaque must be removed, then invasive surgeries, such as arterial bypass angiography, are needed to remove the portion of the blood vessel containing the plaque. However, bypass surgeries are invasive, costly, and risky in elderly patients with comorbidities^12^. Thus, there is a clinical need for a minimally invasive therapy to safely remove plaques, and to date, no therapeutic approaches have provided a cell-specific, mechanical method to selectively decellularize atherosclerotic tissue.

Although a significant portion of the atherosclerotic plaque is fibrotic, the majority of the plaque is cellular, comprised of foam cells, endothelial cells, and SMCs^13^. These cells are the key aggressors in propagating the disease and present a potential target for therapeutic intervention. Electroporation is a biophysical phenomenon in which exogenous pulsed electric fields (PEFs) increase the permeability of the cellular membrane. PEFs generate an ionic current within the tissue, inducing an electric potential across the plasma membrane of cells as charges of opposite polarity accumulate on each side. Once the induced transmembrane potential reaches a critical value (∼0.258 V)^14^, the membrane permeabilizes through the formation of nanoscale defects^15^. If persistent, the cells within a critical electric field threshold die due to loss of homeostasis, a process termed irreversible electroporation (IRE)^16,17^. Pulsed Field Ablation (PFA) is a novel technique that employs high-voltage (1,000 – 3,500 V) microsecond (0.5 – 100 µs) monophasic or biphasic square pulses to induce controlled volumes of IRE within tissues^18^. The high-power, low-energy pulses are applied between electrodes inserted within or adjacent to the diseased tissue to deliver the lethal PEF^19,20^.

PFA has the advantage of cell death being independent of thermal effects, which is paramount for ablating tissues in anatomically sensitive areas where there is concern about damage to nearby critical structures. Proteinaceous structures, such as mature blood vessels^21,22^ and the esophagus^23^, retain their patency even when directly involved in the ablation and when transmural decellularization is observed. Further, PFA does not suffer from the heat sink effect observed with thermal ablation modalities, which may allow targeted tissue around blood vessels to survive treatment. While PFA has not yet been applied directly within arteries for AS treatment, catheter-based PFA has generated considerable attention for the treatment of atrial and ventricular fibrillation, with data from many large clinical trials recently released^24–31^. Cardiac PFA has demonstrated a favorable safety profile, with low rates of vascular stenosis, thrombus formation, or endothelial damage. These findings suggest that PFA can be safely delivered endoluminally without compromising vascular structure or long-term function. PFA is also demonstrated to ablate cells through fibrotic scar tissue^32^ and calcified structures, such as bone^33^, suggesting that it can decellularize fibrotic and calcified plaques. Further, foam cells become larger and more vacuolated than their normal counterparts due to phagocytosis and accumulation of lipids. Due to the increased cell membrane surface area, the induced transmembrane potential is higher for larger cells^34,35^. Further, cell membrane organization significantly affects electroporation outcomes, with higher lipid saturation and cholesterol lowering the threshold for pore formation^36^. Given that AS plaques are rich in disease-driving cell types, the ability of PFA to selectively decellularize plaques while preserving extracellular architecture creates a compelling therapeutic strategy.

We hypothesize that PFA can be delivered through a minimally invasive, endoluminal catheter to induce IRE in the cellular contents of AS that propagate the disease, enabling localized plaque regression or priming for adjunctive therapies (Figure 1A). This cell-specific, nonthermal mechanism will enable controlled, localized decellularization without the risk of heat-induced injury or restenosis commonly observed with stenting or thermal ablation. First, we designed, manufactured, and evaluated a prototype catheter-based system for delivering PFA for endoluminal vascular diseases. We then determined optimal pulsing protocols for lethal electric field coverage within a multi-tissue atherosclerotic vascular disease finite element model. Finally, we developed a 3D fibrotic *in vitro* atherosclerosis microgel model to validate the ability for PFA to selectively eradicate foam cells without damaging the extracellular matrix. These complementary studies establish proof-of-concept and foundational supporting evidence that endoluminal PFA can treat atherosclerotic vascular disease.

**Figure 1.**
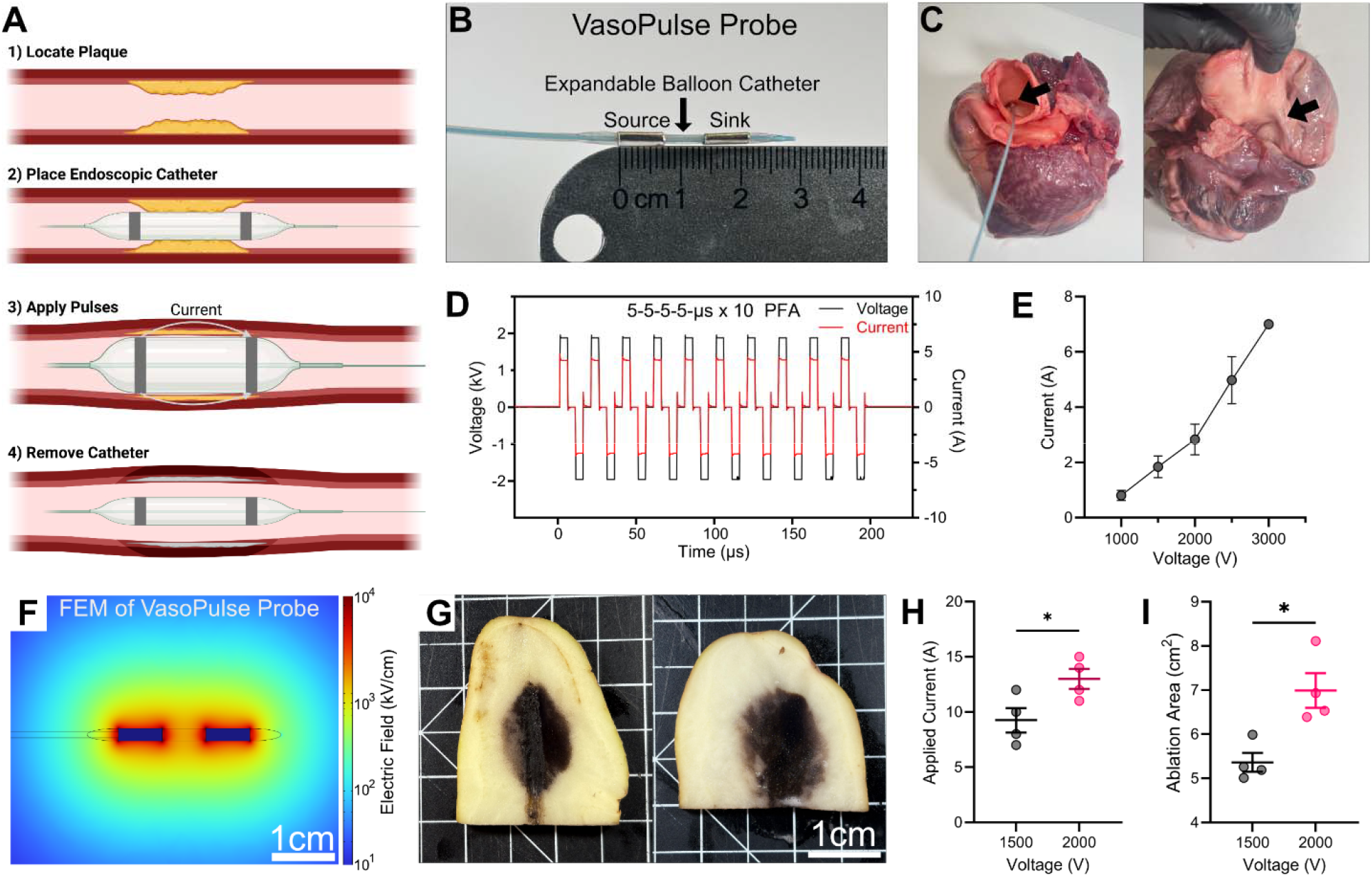
Development and evaluation of a bipolar balloon catheter for endoluminal pulsed field ablation delivery. **A)** Schematic of proposed endoluminal catheter treatment with balloon placement, expansion, and pulsed electric field delivery. **B)** Prototype vascular pulse (VasoPulse) delivery probe fabricated using a flexible conductive strip on top of wires and wrapped around a non-conductive polyethylene terephthalate (PET) endoluminal balloon catheter. **C)** The VasoPulse probe was inserted through the coronary artery of four *ex vivo* porcine hearts for proof-of-concept high-voltage pulse delivery. The arrow points to the coronary artery entrance and the VasoPulse probe within the artery after full insertion. **D)** Representative voltage and current data recorded for a 5-5-5-5-μs x 10 PFA waveform delivered within the coronary artery. **E)** Sequentially increasing electric potentials were delivered to determine maximum deliverable voltage within the prototype VasoPulse probe; mean ± SD; n=4 (n=1 at 3000V). **F)** Simulated electric field distributions of the VasoPulse probe (panel B) within tuber tissue for 2000 V applied between the electrodes. **G)** Representative images of tuber tissue ablation 24 hours after VasoPulse PFA delivery with 1500 V (left) or 2000 V (right). **H)** Measured current delivered during PFA treatment delivery, and **I)** ablation areas measurements; mean ± SEM; Welch’s T-test; * *p <* 0.05; n = 4.

## Results

### Design and evaluation of an endoluminal vascular pulse field ablation catheter

To use PFA for the treatment of endoluminal diseases, an endoscopic catheter must be developed to deliver lethal PEFs within the diseased tissues. A previous single-needle dual-electrode “bipolar” probe used for oncologic PFA delivery was demonstrated to deliver multiple PFA parameters at 2250 V without intraoperative paralytic or cardiac synchronization^37^. In this study, no muscle contractions or cardiac abnormalities were detected, treatment time was 5.0 ± 0.2 minutes, and ablations were visualized post-treatment using ultrasound^37^. We propose that a bipolar design can be adapted for endoluminal vascular PFA delivery to translate the benefits of simplified and rapid PFA delivery. We developed a novel endoluminal pulsing (“VasoPulse”) catheter, which utilizes dual flexible electrodes integrated with a commercial vascular balloon catheter with a 7 mm electrode exposure and 7 mm edge-to-edge spacing (Figure 1B). To evaluate the ability for probe positioning and the feasibility of high-voltage pulse delivery, we positioned the catheter within the left coronary artery of non-perfused *ex vivo* porcine hearts (Figure 1C). After complete insertion of the probe within the coronary artery, we delivered a 5-5-5-5-μs x 10 PFA waveform across the dual electrodes (Figure 1D). To further determine the upper limit of voltage and current deliverable through the prototype VasoPulse probe, we delivered sequentially increasing electric potentials from 1000 V to 3000 V until electrical failure, while recording the voltage and current (Figure 1E). Voltages up to 2500 V were successfully delivered through the VasoPulse probe, but 3000 V was successfully delivered in only 25% (1/4) of the arteries due to electrical arcing. These data suggest that the VasoPulse probe is suitable for PEF delivery within large blood vessels.

We then evaluated the ability of the VasoPulse probe to efficiently deliver energy into tissues and generate contiguous ablations. We utilized *Solanum* Tuberosum vegetal tissue, a common model for assessing PFA in live tissue^38^. We first created a 3D time-dependent finite element model of Tuberosum tissue to simulate the electric field distribution throughout the tissue, considering dynamic electroporation-dependent and thermal-dependent conductivity changes (Figure 1F)^39^. The electric field displays a cylindrical shape previously demonstrated for the dual-electrode type probes^37,40^. Cellular damage within Tuberosum tissue is visible following PFA due to a natural oxidation process of released intracellular enzymes (polyphenol oxidase) and subsequent development of brown-black melanins (Figure 1G)^41^. We previously found that melanin areas correlated with non-viable tissue and were saturated 24 hours after treatment within the Tuberosum tissue^39^, so ablation areas were measured 24 hours after treatment. The applied currents were significantly higher within the Tuberosum tissue than within the *ex vivo* coronary artery for both 1500 and 2000 V. The measured current at 2000 V (13.00 ± 0.91 A) was significantly larger than at 1500 V (9.25 ± 2.22 A; *p <* 0.05) (Figure 1H). Subsequently, the measured ablation areas were also significantly higher at 2000 V (6.99 ± 0.39 cm^2^) than at 1500 V (5.36 ± 0.22 cm^2^; *p <* 0.05) (Figure 1I). The data demonstrate that the VasoPulse probe can efficiently deliver PEF within tissues to generate predictable regions of IRE.

### Simulations of translational endoluminal pulsed field ablation in randomized patient-driven atherosclerotic vascular disease

To demonstrate the translational utility of the VasoPulse probe, we created a time-dependent multi-tissue finite element model of the coronary artery with a plaque embedded between a fibrous cap and the arterial wall (Figure 2A). Joule heating and thermal-dependent conductivity changes were incorporated to quantify safe limitations on voltage deliverable through the VasoPulse probe and calculate the electric field penetration through the tissue. Heterogeneity of plaque composition (i.e., calcified/fibrous structures, lipid, cellular) results in patient-to-patient variability in electrical conductivity. We also incorporated patient-driven geometric and material tissue properties (Table 1), enabling a robust evaluation of the probe and treatment parameters across scenarios with patient-to-patient variability^42,43^. Every replicate within the computational model was defined using the calculated patient electrical conductivities (Figure 2B). The applied current, measured at the last treatment pulse, was significantly different between applied voltages (Figure 2C). The simulated applied currents are comparable to those delivered by the VasoPulse probe within the Tuberosum tissue, suggesting that the probe may be able to successfully deliver them within a viable artery. The simulated electric field increased significantly with applied voltage (Figure 2D) but did not decrease much with depth within a 0.7 mm plaque (Figure 2F). The temperature also increased with increasing applied voltage (Figure 2E), with significant differences between each voltage (Figure 2G). Lastly, we evaluated the percentage coverage of severe and large plaques by an ablative electric field, assuming a conservative lethal threshold of 1000 V/cm, based on previous cardiovascular ablation data using the 5-5-5-5-µs waveform (Figure 2H)^44^. Every plaque size was covered by at least 75% ablative electric field for all voltages. The percent coverage significantly increased with applied voltages across all plaque sizes (*p <* 0.0001), and there were significant differences in percent coverage between the plaque sizes (*p <* 0.05). Together, the plaque coverage and applied current results suggest that 5-5-5-5-μs PFA at 2000 V can potentially achieve sufficient coverage without thermal damage, when delivered through the VasoPulse probe.

**Table 1:**
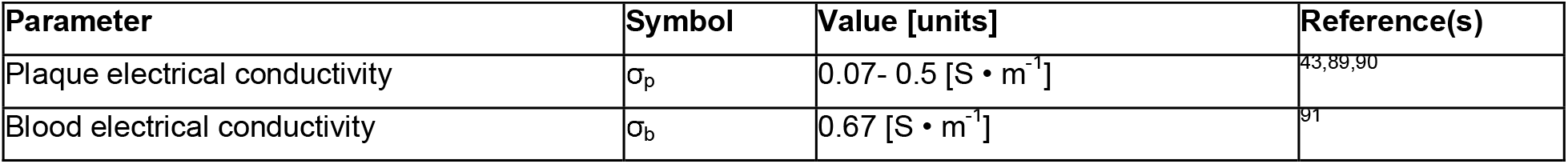
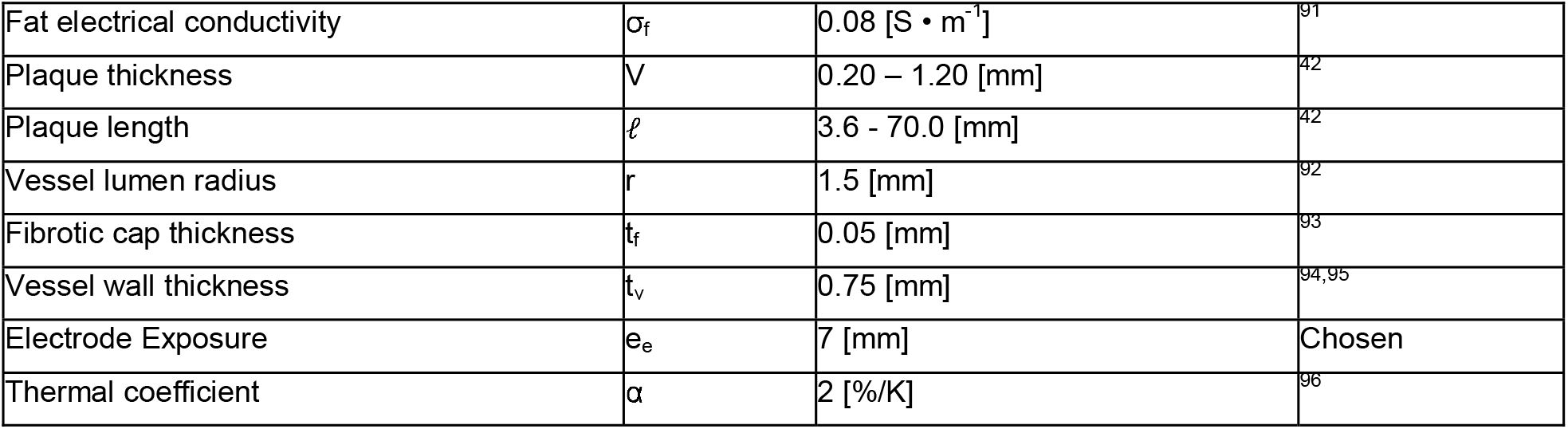
Material and geometric properties for the tissue-level atherosclerosis finite element model.

| Parameter | Symbol | Value [units] | Reference(s) |
| --- | --- | --- | --- |
| Plaque electrical conductivity | $\sigma_p$ | 0.07- 0.5 [$S \cdot m^{-1}$] | 43,89,90 |
| Blood electrical conductivity | $\sigma_b$ | 0.67 [$S \cdot m^{-1}$] | 91 |
| Fat electrical conductivity | $\sigma_f$ | 0.08 [$S \cdot m^{-1}$] | <sup>91</sup> |
| Plaque thickness | $V$ | 0.20 – 1.20 [mm] | <sup>42</sup> |
| Plaque length | $\ell$ | 3.6 - 70.0 [mm] | <sup>42</sup> |
| Vessel lumen radius | $r$ | 1.5 [mm] | <sup>92</sup> |
| Fibrotic cap thickness | $t_f$ | 0.05 [mm] | <sup>93</sup> |
| Vessel wall thickness | $t_v$ | 0.75 [mm] | <sup>94,95</sup> |
| Electrode Exposure | $e_e$ | 7 [mm] | Chosen |
| Thermal coefficient | $\alpha$ | 2 [%/K] | <sup>96</sup> |

**Figure 2.**
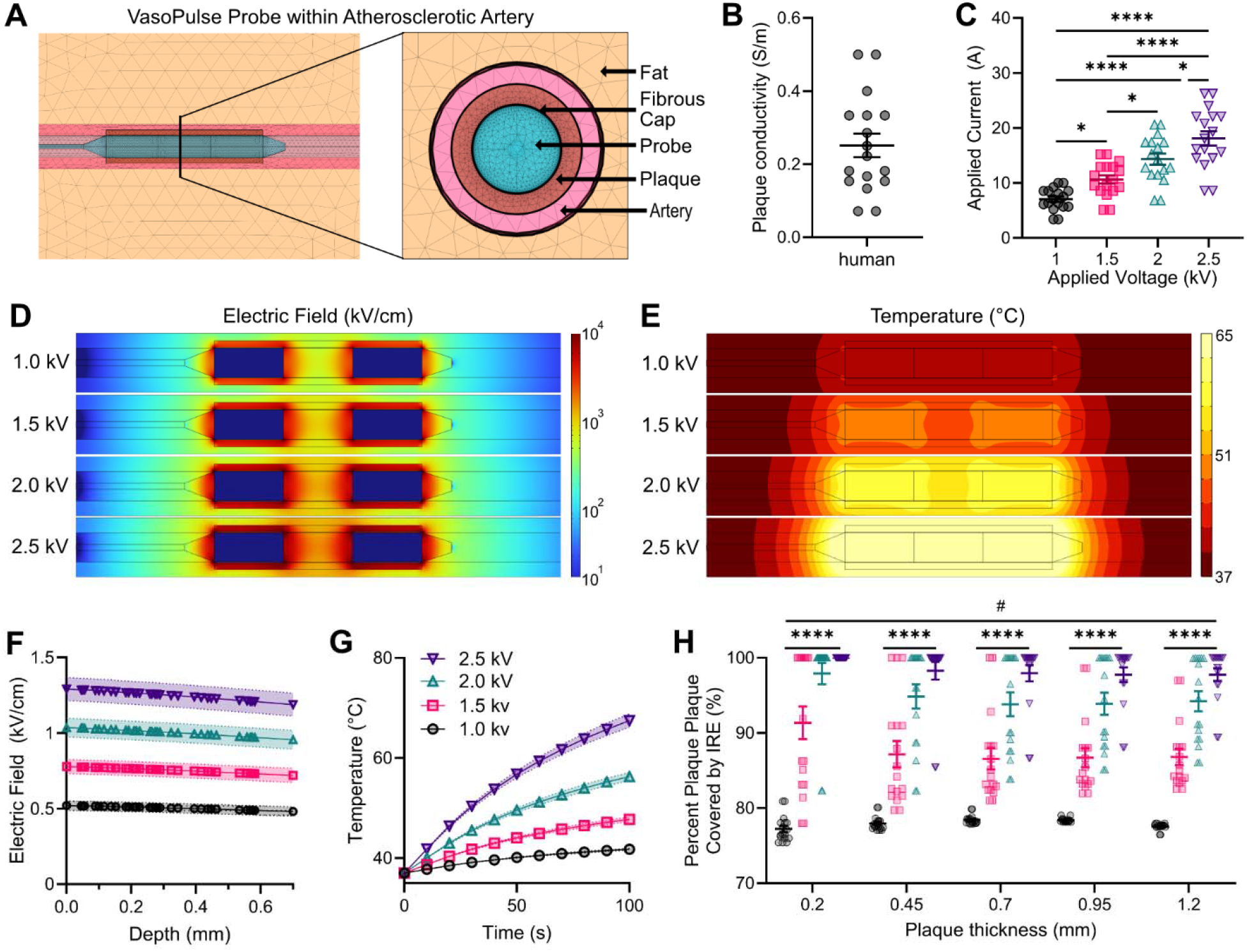
Simulations of pulsed field ablation delivery to endoluminal vascular plaques. **A)** Multi-tissue finite element model of the VasoPulse probe within an atherosclerotic artery (COMSOL™). **B)** Calculated human plaque electrical conductivities from previously characterized impedances^43^; mean ± SEM; n = 17. **C)** Calculated current for a range of applied voltages (1.0-2.5 kV) using patient-derived plaque electrical conductivities; mean ± SEM; One-way ANOVAs with Tukey’s post hoc (* p < 0.05, **** p < 0.0001); n = 17. **D)** Electric field and **E)** Temperature distributions 1 – 2.5 kV; plaque electrical conductivity of 0.25 S/m. **F)** Electric field magnitude off across the plaque depth from the center of the probe radially; mean ± SEM; n = 17. **G)** Temperature at the center point of the plaque over time for different applied voltages (1 – 2.5 kV); mean ± SEM; n = 17. (Panels F & G; repeated One-Way ANOVAs (General Linear Model [GLM]); all electric field groups are significantly different; p < 0.0001; n = 17). **H)** Calculated percent plaque coverage by irreversible electroporation across different plaque thicknesses and applied voltages (1 – 2.5 kV), assuming a lethal threshold of 1,000 V/cm to the 5-5-5-5-μs x 10 pulsed field ablation waveform; mean ± SEM; Two-Way ANOVA (**** p < 0.0001 significance between applied voltages; # p < 0.05 significance between plaque thicknesses); n = 17.

### Analysis of foam cell response to pulse electric fields in a single-cell electroporation model

Electroporation is driven by a change in transmembrane voltage across the cell membrane^14^. The lowest electric field achieved within the plaque at an applied voltage of 2000 V was ∼1000 V/cm (Figure 2D & F); therefore, we investigated the cellular response at electric fields ranging from 1 to 2.5 kV/cm. To examine the response of cells within the plaque, we created a single-cell model of a foam cell with extracellular electric conductivities matching the patient-derived conductivities (Figure 3A). PEFs are indicated to more easily induce TMPs in larger cells^14^, and, with the evagination of oxidized lipids, foam cells grow to larger diameters (∼30 µm) than already big macrophages (∼21 µm)^45^. An induced transmembrane potential (Figure 3B) and subsequent pore formation (Figure 3C) were indicated by the single-cell pore model at electric fields ranging from 1 to 2.5 kV/cm. Pore formation increases the membrane electrical conductivity, which in turn feeds back to negatively affect the induced TMP from the normal current on the membrane. The TMPs were close but still significantly different between all applied electric fields (Figure 3D; *p* < 0.0001). However, the average pore density showed distinct and significant differences between each applied electric field (Figure 3E; *p* < 0.0001). To interrogate electroporation spatially across the cell membrane, we measured the TMP and pore density along the arc length from the front to the back of the cell (Figure 3F). We observed that with higher electric fields, the TMP dropped at the front and back of the cell but increased towards the side (Figure 3G). Consequently, the pore density increased at every spatial location with higher applied electric fields (Figure 3H). These biophysical simulations support that the electric field magnitudes generated by the VasoPulse should induce electroporation within the plaque cellular contents.

**Figure 3.**
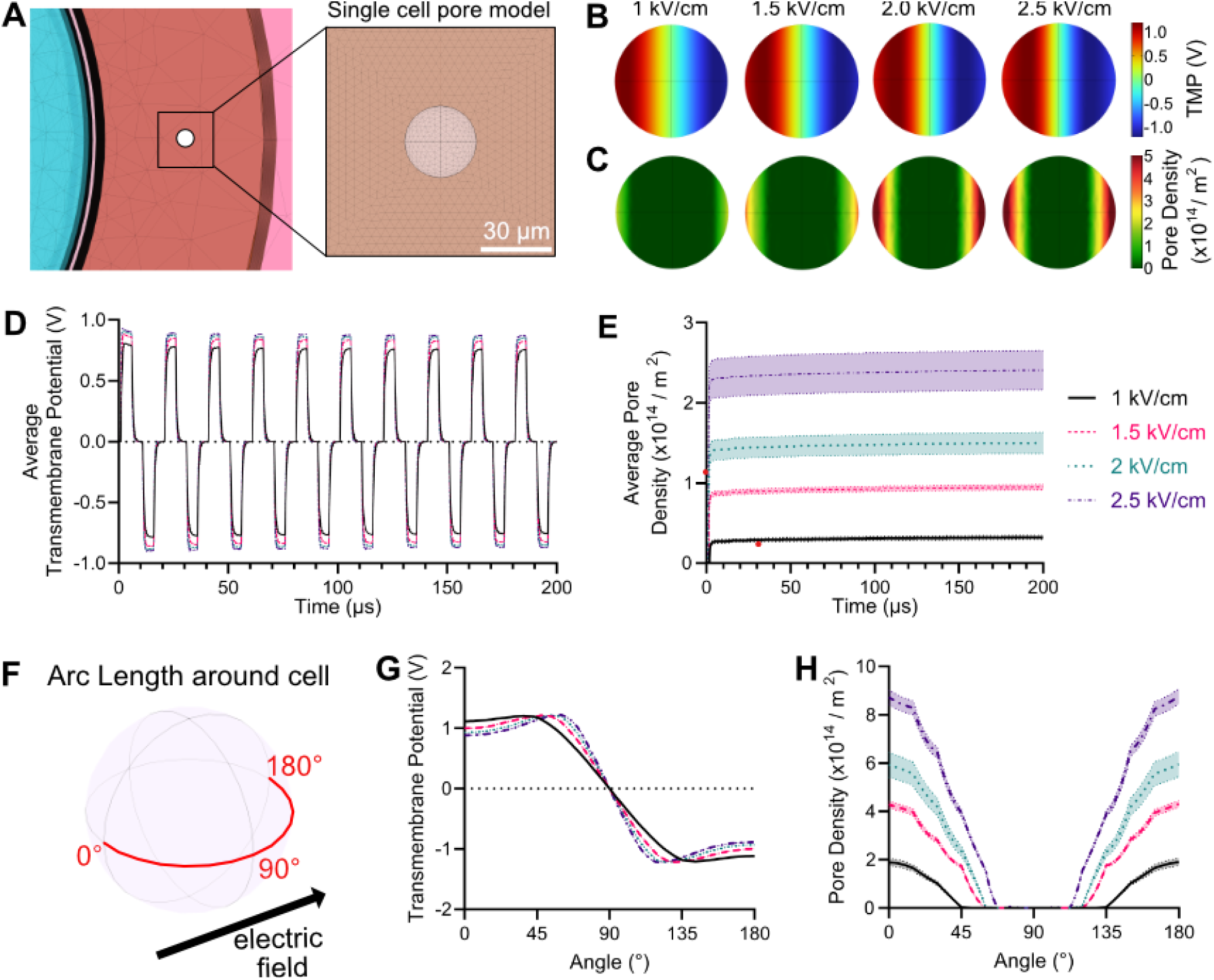
Single foam cell electroporation finite element model. **A)** A single-cell pore model to replicate the electric fields experienced by foam cells within atherosclerotic plaques. **B)** Spatial transmembrane potential and **C)** pore density on the cell membrane in a 1 – 2.5 kV/cm electric field; electric field applied from left to right, and schematic is during the pulse of a 5-5-5-5-μs x 10 PFA waveform. **D)** Average transmembrane potential and **E)** average pore density over the PFA waveform, measured on the left half of the cell due to symmetric but negative TMPs on both sides. **F)** Electric field direction and defined arc length for measuring the **G)** transmembrane potential and the **H)** pore density over the cellular angle and across external electric fields from 1 – 2.5 kV/cm. (Panels D, E, G, and H; repeated One-Way ANOVAs (General Linear Model [GLM]); all electric field groups are significantly different; *p* < 0.0001; n = 17)

### Pulsed field ablation can effectively eradicate foam cells in an *in vitro* atherosclerosis spheroid model

*In vitro* models of AS provide a bridge between computational modeling and *in vivo* validation to determine treatment e□ciency and explore biological mechanisms in a reproducible and high-throughput system^46,47^. Traditional *in vitro* 2D platforms have been used to study AS, but they do not recapitulate the physiological structure observed *in vivo*, have issues with substrate topography and sti□ness^48^, and fail to provide proper pathological compositions of human atherosclerotic plaques^49^. *In vitro* 3D models have been gaining increasing interest for AS research and drug testing, achieving higher biomimicry of cell–cell and cell-ECM interactions^*50,51*^, gene expression, cellular heterogeneity, and microarchitecture of the native tissue^52–54^. Consequently, 3D models improve the predictability of therapeutic toxicity and sensitivity, with significantly higher drug resistance observed in 3D models compared to 2D models^55^. To demonstrate the proof-of-concept treatment of AS using an electroporation-based therapy and determine the PFA parameters that selectively eradicate foam cells, we created an *in vitro* fibrotic AS micro-hydrogel (μGel) model.

For most advanced AS plaques, the majority component is non-cellular fibrosis, composed primarily of collagen^56,57^. Therefore, we encapsulated a high concentration of differentiated THP-1 macrophages and aggregated oxidized LDL within a 3D collagen scaffold (Figure 4A). Oxidized LDL was tagged with Dil fluorescent protein for visualization. Following fabrication, the μGels were treated with a 5-5-5-5-μs x 10 PFA waveform using uniform electric fields from 0 kV/cm (control) to 2 kV/cm (Figure 4B). As the extracellular electrical conductivity affects the induced transmembrane potential and pore electrical conductivity^58–60^, which then ultimately affects IRE thresholds^61^, we also adjusted the conductivity of the pulsing medium to match the measured electrical conductivities for human AS plaques (Figure 4C)^43^. We observed a stark decrease in viability 24 hours after treatment, assessed qualitatively by Calcein AM green fluorescence and validated quantitatively using an XTT metabolic assay (Figure 4D). All treated PFA groups had significant decreases in viability compared to the non-treated control (*p <* 0.0001), with no further differences between the treatment groups. We also observed a non-significant increase in the red-orange fluorescent intensity of the μGel following treatment (Figure 4E). Since both the μGel retained its shape and structure following treatment and the Dil-oxLDL did not diffuse out of the microgel following treatment, this increase in fluorescent intensity may be attributed to the decrease in foam cell viability and subsequent rise in μGel transparency. Together, these data provide proof-of-concept that PFA can selectively decellularize foam cells from a fibrotic tissue and validate the trends observed within the single-cell computational model.

**Figure 4.**
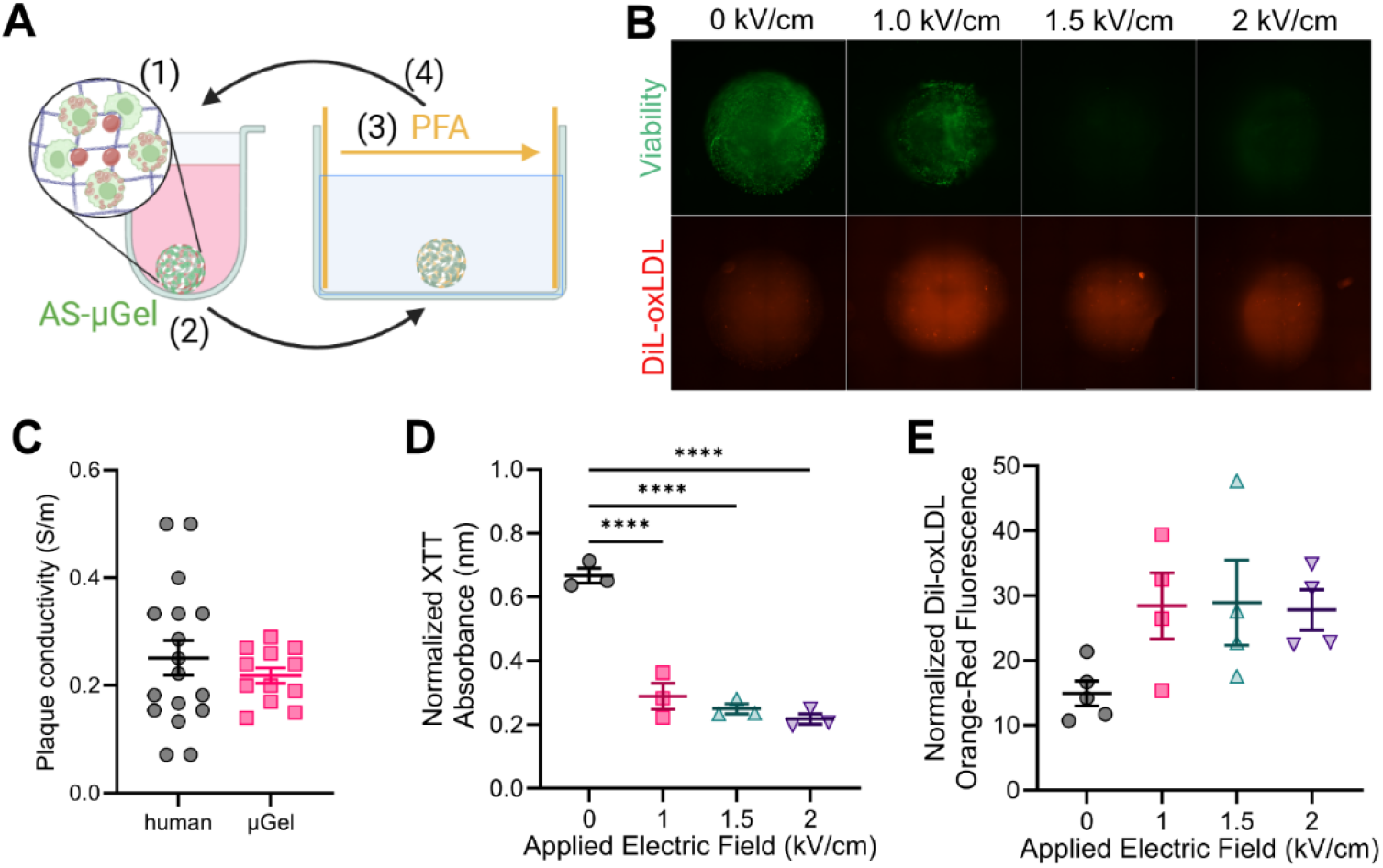
*In vitro* 3D assay to evaluate foam cell death using pulsed field ablation. **A)** *In vitro* atherosclerosis (AS) micro-hydrogel (μGel) assay to assess foam cell treatment response to pulsed field ablation (PFA). (1) Macrophages were combined with Dil-oxLDL and collagen hydrogel to form μGels, (2) then moved to a 4-well rectangular plate with low-conductivity buffer to (3) generate ablative uniform electric fields via parallel plate electrodes. (4) The μGels were immediately moved back into a U-bottom well for follow-up analyses. **B)** Live (green) and DiL-oxLDL (orange-red) imaging of μGel 24 hours after PFA; scale bar is 1 mm. **C)** Calculated human plaque electrical conductivity and measured μGel electrical conductivity; mean ± SEM; Welch’s T-test; n ≥ 12. Normalized absorbance for the XTT viability assay at 24 hours post-treatment across different electric fields; One-Way ANOVA with Tukey’s post-test and correction; mean ± SEM; n=3. **E)** Normalized DiL-oxLDL red-orange fluorescent intensity between electric fields; mean ± SEM; One-way ANOVAs with Tukey’s post-test and correction; n ≥ 4.

## Discussion

Atherosclerotic vascular disease remains a leading cause of morbidity and mortality worldwide, and current treatment strategies such as angioplasty, stenting, and atherectomy are limited by restenosis, thrombosis, and incomplete long-term efficacy. Thermal ablation modalities are not suitable for vascular applications due to collateral tissue damage and the heat sink effect near flowing blood. PFA has emerged as a safe and effective tool in cardiac electrophysiology, particularly in pulmonary vein isolation, where it demonstrates minimal risk of vascular stenosis or structural injury. Building on this safety profile, we present the first proof-of-concept for endoluminal PFA to treat atherosclerotic plaques. We establish feasibility through a combination of prototype probe stress-testing within ex *vivo* porcine coronary arteries, generation of IRE with the prototype probe, multi-tissue computational simulations that quantify IRE coverage within severe patient-derived plaques, and *in vitro* validation of foam cell ablation without compromising extracellular matrix integrity. These findings provide the foundation for developing endovascular PFA as a novel therapeutic strategy for vascular disease.

Unlike drug therapies that require lengthy and expensive development, production, and quality control, electroporation-based therapies are hardware-driven, making them inherently more scalable. Further, electroporation-based therapies are not limited by blood-tissue distribution, receptor expression, or cellular uptake. Irreversible electroporation can be precisely induced in virtually any cell type, producing well-defined ablation zones with sharp and predictable boundaries, even in heterogeneous tissues such as vascular plaques. This makes the therapy suitable for repeated applications at the same or different sites, without concerns about cumulative toxicity, maximum dosing, or systemic interactions. As a result, PFA offers a flexible, durable strategy for managing complex or multifocal atherosclerotic disease. Since no systemic drug is administered, there is also minimal risk of off-target toxicity to the heart, liver, kidneys, or other organs. This is particularly valuable for high-risk patients with multiple comorbidities, for whom systemic anti-inflammatory or lipid-lowering drugs may carry undesirable side effects^62,63^. Therefore, unlike many pharmacologic agents that lose efficacy over time due to receptor downregulation or adaptive resistance, there is no known mechanism for cellular resistance to IRE. However, PFA does not preclude the concomitant use of therapeutics, as the ability of small and large molecule drugs to distribute into the diseased tissue may increase. Adiponectin is a well-known circulating adipokine with antidiabetic and antiatherogenic properties that can bind to and neutralize oxidized lipids^64^. Electroporation may kill the cellular contents of the atherosclerotic plaque while facilitating the delivery of therapeutics that target the non-cellular contents.

We chose to use a biphasic PFA waveform to reduce muscle contractions associated with conventional monophasic IRE, which can cause an involuntary shift in the electrode locations and incomplete ablation of the target region^17,65–67^. Our group has also demonstrated that PFA is well-tolerated in five awake veterinary equine patients without general anesthesia, validating the reduction in pain and muscle contractions^68^. While shorter pulse widths can mitigate muscle contraction, the reduced membrane charging with shorter biphasic waveforms increases the EF required to induce IRE^69,70^. We found that 5-μs pulse widths could completely eradicate foam cells and potentially cover more than 95% of severe and large AS plaques with an ablative field, so future studies should investigate shorter pulse widths to retain full plaque electroporation while minimizing deleterious thermal and neuromuscular effects^71,72^. The PFA waveform is highly parameterizable, and increasing the inter-pulse delay (d_2_) and shortening the interphase delay (d1) can further reduce or abrogate neuromuscular excitation^73–76^, without affecting the final ablation volume^39^. Further, bursts can be delivered slowly to facilitate thermal heat diffusion or convection^77,78^ or deliver fewer bursts after electroporation is saturated to minimize excess energy delivered^69,79^.

Along with waveform optimization, the probe design and procedure would need to be optimized. We expect that endoluminal PFA would follow well-established catheter delivery techniques and is expected to have a similar cost and time profile to stent placement, enabling rapid integration into existing clinical workflows. Moreover, we demonstrate that the treatment zone can be accurately modeled using established computational methods, enabling personalized therapy planning based on vessel size, plaque burden, and conductivity profiles. The prototype VasoPulse probe was developed with similar geometric parameters to the bipolar probe, and the maximum voltages presented here closely match those previously observed^40^. For the prototype VasoPulse probe, multiple designs with varying lengths would be required, depending on the target treatment length. However, we observed that the area of the plaques that fell below the ablative field was in the middle between the electrodes, so it would be advantageous to develop a probe that accounts for patient-to-patient variability in plaque length and the drop near the center by utilizing multiple electrodes. Single electrode catheters coupled with a distant ground and shorter pulse widths have been utilized to create uniform lesions^80^. The prototype VasoPulse design utilizes a balloon catheter; however, multielectrode arrays can be integrated with stent-like constructions to both treat various plaque lengths and facilitate blood flow during treatment, reducing ischemic time and increasing thermal heat convection. Future work should validate the use of PFA for AS and optimize the probe design with *ex vivo* and *in vivo* animal models of atherosclerosis^81^. Further, the atherosclerotic tissue mimic developed here can be expanded to incorporate other disease-driving cell lines in a self-assembling organoid. This would be beneficial for high-throughput analysis of PFA parameters or biological responses, as there are many well-established techniques to obtain qualitative and quantitative data^54^, including gene expression^82,83^, metabolism^84^, cellular motility di□erentiation^85–87^, and polarity of cells^88^.

## Conclusion

Endoluminal pulsed field ablation represents a promising nonthermal approach for treating atherosclerotic vascular disease. Unlike existing therapies, it selectively targets the cellular components of plaque while preserving the underlying vessel scaffold, suggesting potential to reduce restenosis and maintain long-term vessel patency. Clinically, such a technology could be delivered through standard endovascular workflows alongside angioplasty or stenting, or as a stand-alone therapy in patients with diffuse or treatment-resistant disease. This could allow for earlier intervention in patients with AS, reduce the need for surgical bypass, and improve outcomes in patients at high surgical risk. Additionally, this approach opens a new therapeutic paradigm in which plaques can be sensitized or cytoreduced before drug delivery or stenting, thereby meeting a critical gap in the treatment of intermediate or advanced atherosclerotic disease. These features position pulsed field ablation as a potential alternative to conventional interventions, warranting further translational studies in large animal models and ultimately in patients.

## Methods

### Endoluminal bipolar electrode balloon catheter pulse field ablation probe fabrication and evaluation

The prototype endoluminal bipolar balloon catheter (“VasoPulse”) probe was fabricated from a non-conductive 30-mm cylindrical balloon catheter (Stryker, Kalamazoo, USA), with two 7-mm flexible conductive electrodes spaced 7 mm apart. Thin insulated wires were attached under the electrodes and fed down the shaft to allow for electrical connections to the pulsed electric field generator. Pulsed electric fields were delivered using a custom high-voltage pulsed electric field generator (Vitave Inc., OmniPorator, Czech Republic). The applied voltages and currents were recorded using a WaveSurfer 5 GHz oscilloscope (Teledyne LeCroy, 4024HD) equipped with a 1000× attenuated high-voltage probe (Siglent, DPB5700) and a 10× attenuated current probe (Pearson Electronics, 3972). To stress-test the prototype VasoPulse probe within a realistic geometry, we thawed whole frozen porcine hearts (<1 month frozen) to room temperature (20°C, measured). The hearts were originally obtained at the University of Georgia Meat Science Technology Center (Athens, Georgia, USA) from female Yorkshire-Landrace pigs (1-2 years old, estimated 200kg at euthanasia). The VasoPulse probe was inserted down the aorta and positioned entirely within the left coronary artery, ensuring good electrical contact with the vessel walls. To evaluate the maximum voltage, 100 pulse trains of the 5-5-5-5-μs x 10 waveform were applied at a rate of 1 Hz, with the applied voltage increased after every successful voltage delivery until failure by electrical shorting.

### Ablation generation using the endoluminal pulsed field ablation probe

*Solanum* Tuberosum (Russet potatoes) were cut in half perpendicular to the longest axis. A small guide hole was generated, and the VasoPulse probe was fed ∼ 2 cm deep into the tissue. Pulsed electric fields were generated as described above using the custom high-voltage generator, with the voltage and current recorded. The tissues were maintained for 24 hours at room temperature to allow for melanin to develop. After 24 hours, we cut down the guide hole and then imaged the ablation on a grid cutting mat using a digital camera with a metric ruler. A custom holder was used to maintain 16.5 cm from the surface for every image. The images were then imported into ImageJ (NIH) for analysis. The dark melanin region for each potato was measured using the freehand selection tool. The pixel value for the length of the ruler was used to convert the measured pixel areas from ImageJ into centimeters squared.

### Multi-tissue atherosclerotic vascular disease computational model

The 3D, time-dependent multi-tissue model was developed within COMSOL™ Multiphysics 6.2, with the material and geometric properties given in Table 1. The electric field distribution and Joule heating within the tissues were modeled using the AC/DC and heat transfer models, respectively. Current conservation is modeled within COMSOL using a modified Laplace equation under a quasi-static approximation. One electrode was defined as the voltage source, and the other was defined as the voltage sink. We modified the heat generation within solids to solve the bioheat equation with the addition of Joule heating from the applied electric field:

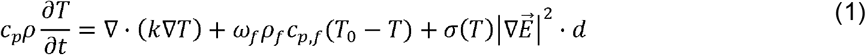

where cp is the tissue specific heat, *ρ* is the tissue density, *k* is the tissue thermal conductivity, *ω* is the tissue perfusion frequency, and *d* is the duty cycle for the applied burst. The blood around the stent was considered stagnant; therefore, tissue perfusion was only considered in the fat domain surrounding the vessel. For initial modeling purposes, we assumed that tissue electrical conductivity does not change with electroporation effects. Instead, electric conductivity is only a function of local temperature. The local electrical conductivity of each tissue is then dynamically updated by the increase in local temperature:

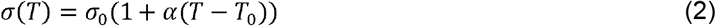

where *σ*_0_ is the initial electrical conductivity of the tissue at body temperature, *T*_0_. The thermal coefficient, *α*, is the percent increase in electric conductivity from an increase in temperature. The overall output of this finite element model is the electric field and temperature distributions in the hydrogel over time.

### Single foam cell finite element model

The 3D, time-dependent single-cell model was developed within COMSOL™ Multiphysics 6.2.^97,98^ using the material and geometric properties defined in Table 2. Current conservation was modeled as:

**Table 2:**
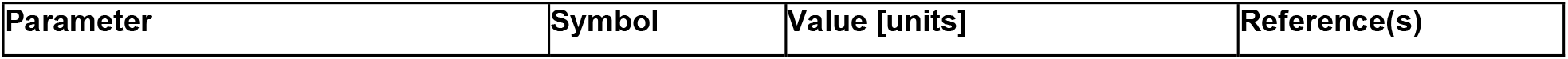
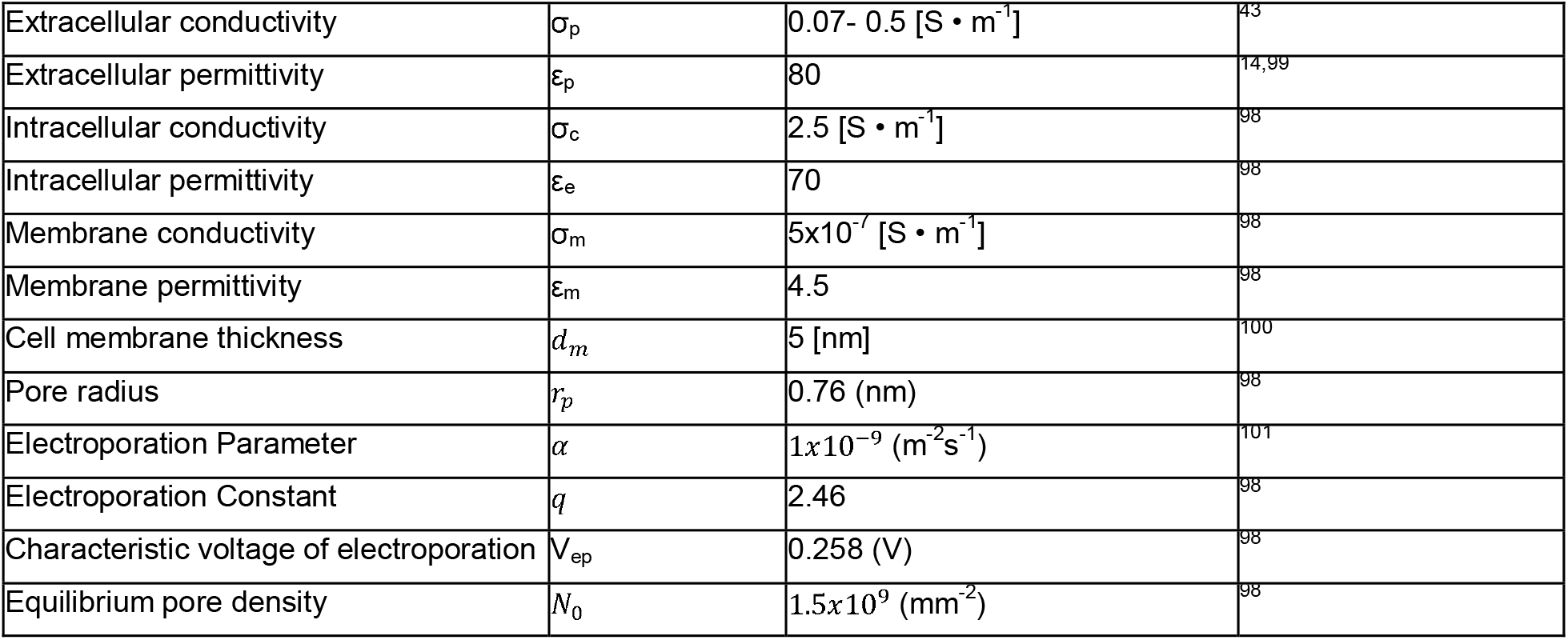
Material and geometric properties for the single-cell atherosclerosis finite element model.

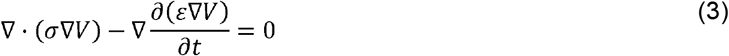

Where *α* is the material electrical conductivity and *ε* is the material electrical permittivity. To model the membrane transmembrane voltage, we defined a distributed impedance boundary condition on all the thin layer cell membrane surfaces:

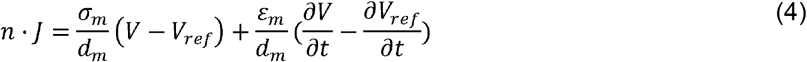

Where *n* is the unit vector normal to the cell membrane surface, *J* is the current flux, *σ*_*m*_ is the membrane electrical conductivity, *dm* is the thickness of the membrane *V* is the electric potential outside of the membrane, *Vref* is the voltage inside the membrane, and *ε*_*m*_ is the membrane electrical permittivity. Here the induced transmembrane potential is calculated at every point on the cell membrane surface:

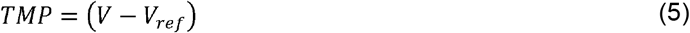

Pore density was calculated using the standard electroporation pore model at every point on the surface using the induced TMP:

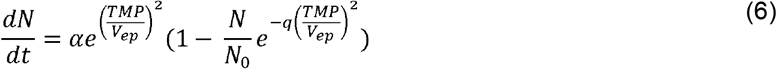

Where *N* is the pore density on the membrane surface, *α* and *q* are fit parameters describing the electroporation process, *V*_*ep*_ is the threshold voltage for electroporation, and *N*_0_ is the equilibrium pore density. Pore formation alters membrane conductivity, which is described as:

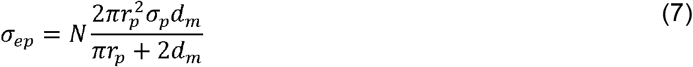

Where *σ*_*ep*_ is the electroporated membrane conductivity at that spot on the surface, *r*_*p*_ is the pore radius, and *σ* _*p*_ is the pore electrical conductivity, determined as:

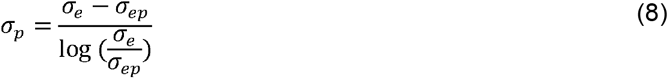

Thus, the dynamic electrical conductivity of a specific location in the cellular membrane is evaluated as:

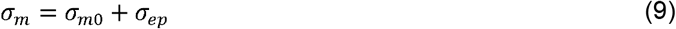

Where *σ* _*mo*_ is the initial membrane conductivity.

### Cell Culture

THP-1 human monocytes (ATCC, TIB-202) were cultured in RPMI-1640 medium (ThermoFisher, 11875093) supplemented with 10% (v/v) fetal bovine serum (Fisher Scientific, FB12999102) and 1% (v/v) 10,000 U/ml penicillin-streptomycin (Gibco, 16140122). The cells were maintained at 37ºC and 5% CO2 in a regulated, humidified incubator (Fisher Scientific, 13-998-086). THP-1s were maintained in cell suspension between 1×10^6^ and 2×10^6^ cells/ml and utilized between passages 2 and 3.

### Collagen-encapsulated foam cell microgel experiments

THP-1s were resuspended in supplemented cell culture media at 20×10^6^ cells/ml with the addition of 50 ng/mL phorbol 12-myristate 13-acetate (PMA, Millipore Sigma, P1585), 1 ng/mL lipopolysaccharide (LPS, Millipore Sigma, L2630), and 50 μg/ml of red-orange fluorescent DiI dye oxidized low-density lipoprotein (Dil-oxLDL, ThermoFisher, L34357) from human plasma^102,103^. To mimic the extracellular environment of AS plaques, neutralized collagen scaffolds were prepared on ice; 10 mg/mL collagen was mixed with 10x DMEM (10% of collagen volume) and with 1 M NaOH (2% of collagen volume)^69,70,104^. An equal volume of the neutralized collagen was combined with the THP-1 suspension to produce a collagen-cell mixture with 10×10^6^ cells/ml and 5 mg/ml collagen. 5 μl of the mixture (50,000 cells per microgel) was pipetted into a V-bottom plate (ThermoFisher, 277143) on ice. After placing all the mixtures, the plate was centrifuged at 300 × g for 1 minute to center the hydrogels. The plate was then incubated for 30 minutes to polymerize the hydrogels, and 100 μl of supplemented cell culture media with PMA and LPS was added to each well. The plates were then incubated for 24 hours. The µGels with 100 µl of cell culture media were individually removed and placed in a rectangular 4-well plate (Ibidi) containing 400 μl of osmotically- and pH-balanced low-conductivity electroporation buffer (0.01 S/m, measured)^105^. Pulsed electric fields were delivered through custom flat plate electrodes with an 8.5 mm spacing^97^, using the custom pulsed electric field generator (Vitave Inc., OmniPorator, Czech Republic). 100 bursts of 10 x 5-5-5-5 μs pulse trains were applied at a rate of 1 Hz, with the applied voltage adjusted to deliver 1000 - 2000 V/cm. The applied voltages and currents were recorded using a WaveSurfer 5 GHz oscilloscope (Teledyne LeCroy, 4024HD) equipped with a 1000× attenuated high-voltage probe (Siglent, DPB5700) and a 10× attenuated current probe (Pearson Electronics, 3972). Controls were similarly placed within the 4-well plate with low-conductivity buffer, but without pulsing. After treatments, the µGels were moved to a BIOFLOAT™ 96-well Ultra-Low Adherent plate (faCellitate, F202003) with 200 μl of fresh supplemented cell culture media.

### Viability and Fluorescent Intensity Imaging

24 hours after treatment, the cell culture media was replaced with a live/dead stain consisting of 0.4 μl Invitrogen™ CellTrace™ Calcein Green, AM (ThermoFisher, C3100MP) in 200 μl of phosphate buffered saline (PBS, ThermoFisher, 10010023), then imaged after 30 minutes at 10x using a Leica DMI8 fluorescent microscope. Normalized fluorescent intensity was quantified within ImageJ Fiji (NIH) by separating the Green and Red-Orange image channels, then subtracting the intensity measured within a region of interest drawn around the µGel by the background intensity.

### Quantitative XTT metabolic assay

The XTT assay was performed according to the manufacturer’s instructions. 6 hours post-treatment, the cell culture media was replaced with 100 μl of fresh supplemented cell culture media and 50μl of the XTT-activated assay reagent (1:50 ratio of Activation Reagent PMS to XTT solution). After incubating for 18 hours, the supernatant within each well was then transferred to a flat-bottom 96-well plate, and absorbance was recorded at 475 and 660 nm (Biotech Synergy HT). The measured absorbance at 475 nm was normalized using the readings at 660 nm.

### Statistical Analyses

Statistics are defined as mean ± SEM within the text and figures. Statistical analyses were performed using GraphPad Prism 10.5.0 (GraphPad Software, Boston), with the specific tests used outlined within the figure legends. Post-experimental power analyses were performed using G*Power 3.1 (Heinrich Heine Universität, Düsseldorf)^106^.

## Acknowledgements and Funding

We would like to acknowledge the Margaret P. and John H. Weitnauer Jr. endowed chair at Georgia Tech for funding support; Harl Ryan Crowe and the University of Georgia Edgar L. Rhodes Center for Animal & Dairy Science for their assistance in cardiac tissue procurement.

## Data Availability Statements

The data that support the findings of this study are available within the article and its supplementary material. Further requests can be made to the corresponding author.

